# A high-continuity and annotated tomato reference genome

**DOI:** 10.1101/2021.05.04.441887

**Authors:** Xiao Su, Baoan Wang, Xiaolin Geng, Yuefan Du, Qinqin Yang, Bin Liang, Ge Meng, Qiang Gao, Sanwen Huang, Wencai Yang, Yingfang Zhu, Tao Lin

## Abstract

Genetic and functional genomics studies require a high-quality genome assembly. Tomato (*Solanum lycopersicum*), an important horticultural crop, is an ideal model species for the study of fruit development. Here, we assembled an updated reference genome of *S. lycopersicum* cv. Heinz 1706 that was 799.09 Mb in length, containing 34,384 predicted protein-coding genes and 65.66% repetitive sequences. By comparing the genomes of *S. lycopersicum* and *S. pimpinellifolium* LA2093, we found a large number of genomic fragments probably associated with human selection, which may have had crucial roles in the domestication of tomato. Our results offer opportunities for understanding the evolution of the tomato genome and will facilitate the study of genetic mechanisms in tomato biology. Information for the assembled genome SLT1.0 was deposited both into the Genome Warehouse (GWH) database (https://bigd.big.ac.cn/gwh/) in the BIG Data Center under Accession Number GWHBAUD00000000.

## Introduction

Tomato (*Solanum lycopersicum*) is an important model plant for scientific researches on fruit development and quality(Meissner et al., 1997). The tomato cultivation area has increased by ~1 million hectares over the past decade, and the yield has increased from 155 million tons to 181 million tons (http://www.fao.org). As a nutritious vegetable that contributes to the human diet, tomato is reported to contain more health-promoting compounds such as lycopene than some other popular fruits. These compounds lower risk of cancer and maintain human health(Giovannucci, 1999). Tomato was originally found mainly in the Andean mountains of South America. Its fruit weight and quality differ markedly among different horticultural groups, and wild tomatoes have smaller seeds and lower yields than cultivars.

A draft genome of the tomato cultivar Heinz 1706 produced using shotgun sequencing technology was released in 2012 (The Tomato Genome Consortium, 2012) and widely used as a reference genome for scientific researches. However, the fragmented nature of this genome and the resulting incomplete gene models could hindered the discovery and functional analysis of important genes. The completeness, accuracy, and contiguity of genome assemblies depend mainly on sequencing technology and assembly strategy. In the current genomic era, single-molecule real-time (SMRT) sequencing technology and new assembly pipelines have remarkably improved the quality of genome assemblies such as those of rice(Du et al., 2017), cucumber(Li et al., 2019), and tomato(Hosmani et al., 2019). Although these genome assemblies have accelerated some scientific researches, such as QTL mapping and transcriptome analysis, higher continuous and complete genome sequences are required for identification of large structural variations and gene mining.

In this study, we generated a highly continuous and complete genome sequence of Heinz 1706 (version SLT1.0) that contains many fewer gaps and unplaced contigs and demonstrates better assembly of repetitive regions. By comparing the genomes of *S. lycopersicum* and *S. pimpinellifolium* LA2093, we found a large number of genomic fragments that appear to be likely involved in domestication. Our work offers new opportunities for understanding the evolutionary history of the tomato genome and the genetic mechanisms that underlie complex traits in tomato breeding.

## Results and Discussion

### High-quality genome assembly

We assembled a highly continuous and complete genome sequence of Heinz 1706 using an integrated genome sequencing approach that combined 131.78 Gb (168.52×) of SMRT data, 226.97 Gb (290.24×) of BioNano data, 140.52 Gb (179.70×) of Hi-C data, and 50.93 Gb (61.53×) of Illumina short-read data (Supp Table 1). The PacBio long reads with an N50 read length of 32.82 kb were assembled with CANU software(Koren et al., 2017), generating a 875.21-Mb genome with a contig N50 of 17.83 Mb (Table 1). To reduce fragmentation and fill in gaps, BioNano data and Hi-C data were used to assist with scaffold construction using Aigner and Assembler(Shelton et al., 2015), HERA(Du et al., 2019), and Juicer(Durand et al., 2016) software. A Hi-C-based physical heatmap comprising 12 groups was generated (Supp. Figure 1) and used to create 12 pseudo-chromosomes that anchor ~790.59 Mb of the genome and harbor 97.61% (33,562) of the predicted protein-coding genes. The genome assembly was polished with Illumina short reads for error homozygous SNPs or indels using Pilon software(Walker et al., 2014). As a result, we generated a 799.09-Mb genome assembly, SLT1.0 (Figure 1 and Table 1).

**Table 1:** Genome assembly and annotation of SLT1.0.

|  | SLT1.0 | SL4.0 | SL3.0 |
| --- | --- | --- | --- |
| Genome assembly (Mb) | 799.09 | 782.52 | 828.08 |
| Non-N bases | 797,955,212 | 782,475,302 | 746,357,470 |
| Number of gaps* | 99 | 259 | 18,271 |
| Number of total contigs | 1,615 | 504 | - |
| Longest contig length (Mb) | 47.16 | 26.29 | - |
| N50 of contigs (Mb) | 17.83 | 6.01 | - |
| Number of unplaced contigs | 112 | 176 | 4,374 |
| Unplaced contigs sequence length (Mb) | 8.50 | 9.64 | 20.85 |
| Number of genes | 34,384 | 34,075 | 35,768 |
| Percentage of gene length in genome (%) | 16.21 | 15.56 | 17.33 |
| Mean gene length (bp) | 3,766.53 | 3,572.44 | 4,011.09 |
| Gene density (per Mb) | 43.03 | 43.55 | 43.19 |
| Mean coding sequence length (bp) | 223.02 | 228.01 | 219.97 |
| Mean exon length (bp) | 310.11 | 275.03 | 308.36 |
| Mean intron length (bp) | 270.41 | 606.69 | 632.38 |
| Masked repeat sequence length (Mb) | 558.49 | 546.95 | 507.14 |
| Repeats percentage of genome size (%) | 69.89 | 69.90 | 61.24 |

\*Not included unplaced contigs

**Figure 1:**
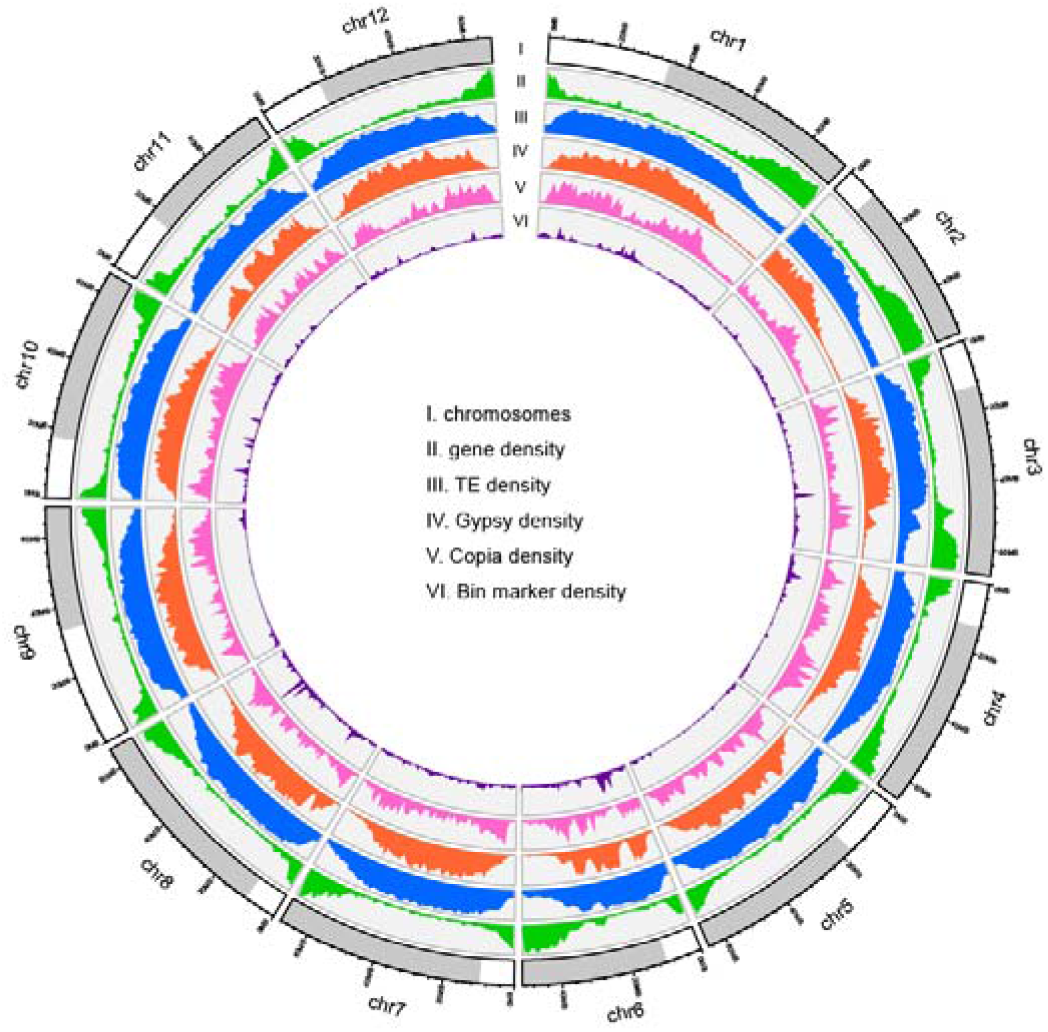
Genomic landscape and structural variants of *S. lycopersicum* cv. Heinz 1706. (i) Ideogram of the 12 chromosomes with scale in Mb. (ii) Gene density (number of genes per Mb). (iii) Repeat content (% nucleotides per Mb). (iv) *Gypsy* content (% nucleotides per Mb). (v) *Copia* content (% nucleotides per Mb). (vi) Bin marker content (% nucleotides per Mb)

The conserved genes from the Benchmarking Universal Single-Copy Orthologs (BUSCO) gene set(Simao et al., 2015) were used to gauge the accuracy and completeness of the SLT1.0 assembly. The results showed that the SLT1.0 assembly contained 97.70% complete genes and 0.30% fragmented genes. The value of the LTR Assembly Index (LAI) was 12.41, which was consistent with that of the previously released SL4.0 tomato reference genome (LAI 12.54). More than 99.88% of the genome assembly had greater than one-fold coverage with Illumina short reads. All these evidences demonstrated the high continuity and completeness of the SLT1.0 genome assembly.

### High-quality genome annotation

Except for *ab initio* prediction and protein-homology-based prediction, we also used transcriptome data, including the bulked RNA-seq data with a mapping rate of 99.73%, and previously-released RNA-seq data from various tissues (The Tomato Genome Consortium, 2012) with a mapping rate of 97.97%, to facilitate gene annotation of the assembled genome. In total, we predicted 34,384 protein-coding genes with an average length of 3,766.53 bp and 6.55 exons per gene in the SLT1.0 genome (Table 1 and Supp Table 2). Gene completeness was estimated to be 98.20% based on the BUSCO gene set(Simao et al., 2015), and the protein-coding genes were unevenly distributed along the chromosomes (Figure 1). Comparative analysis showed that 234 genes in the SLT1.0 genome corresponded to 488 genes in the SL4.0 genome (Supp Table 3). Gene collinearity analysis identified 33 collinear gene blocks between the SLT1.0 and SL4.0 genomes, harboring 28,892 (84.03%) and 28,389 (83.30%) homologous genes, respectively (Figure 2A). Some unplaced contigs in the SL4.0 genome were successfully assigned to chromosomes in the SLT1.0 genome. These results highlight the high accuracy and completeness of the SLT1.0 genome assembly and gene models.

**Figure 2:**
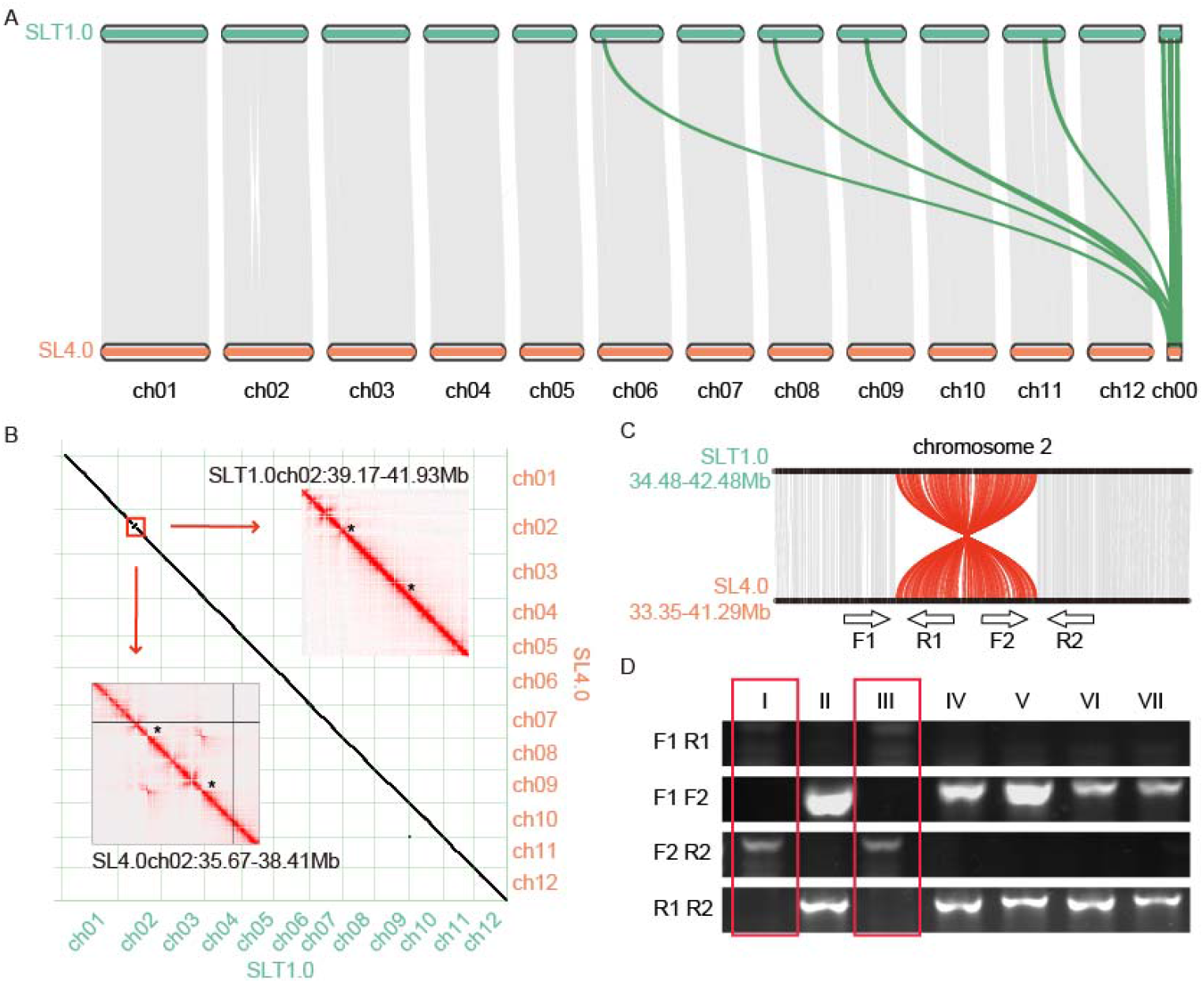
Alignment between the Heinz 1706 SLT1.0 and SL4.0 genomes. **A** Genome collinearity analysis showed that four scaffolds from SL4.0 are placed on chromosomes of the SLT1.0 genome and that there is an inversion on chromosome 2. **B** The color intensity of the Hi-C heatmap represents the number of links between two 25-kb windows. The presence of an inversion is supported by high-density contacts indicated by two asterisks in the Hi-C heatmap generated from SL4.0 Hi-C reads (lower left), whereas no corresponding contact is found in the SLT1.0 Hi-C heatmap (upper right). **C** The inversion shown in red on chromosome 2. F1, R1, F2, and R2 are primers around the break points. **D** Seven Heinz 1706 individuals were identified, two of which (I, III) had inversions

A comprehensive analysis of the genome sequences identified 965 collinear chromosomal blocks between the SLT1.0 and SL4.0 genomes. These blocks contained 32,922 and 32,554 genes, accounting for 95.75% and 95.54% of the SLT1.0 and SL4.0 genomes, respectively. However, we detected a 2.76 Mb inversion from 39.17 to 41.93 Mb on chromosome 2 of the SLT1.0 genome (Figure 2B). The continuous interaction signals on the Hi-C heatmap, as well as PCR and Sanger sequencing, showed that this region was not misassembled (Figure 2B, C and Supp Table 4). This result indicated that heterozygous variation may exist in the previously reported Heinz 1706 accession.

### Transposable element analysis

A total of 524.84 Mb of repetitive sequences were identified, accounting for 65.66% of the SLT1.0 genome assembly, which was similar to that reported in the SL4.0 genome (508.89 Mb, 65.03%) (Supp Table 5). Among these repetitive sequences, long terminal repeats (LTRs) were the predominant TE family, covering 50.25% (401.60 Mb) of the genome. *Gypsy*-type LTRs (344.52 Mb) were the most common subfamily and six times more abundant than *Copia*-type LTRs (50.09 Mb). We used a combination of methods, including LTR-FINDER(Xu et al., 2007), LTR-Harvest(Ellinghaus et al., 2008), and LTR-Retriever(Ou et al., 2018), to identify intact LTRs. A total of 3,220 LTRs were detected in the SLT1.0 genome assembly, including 1,553 *Gypsy*-type LTRs and 1,346 *Copia*-type LTRs. The estimated insertion time of the LTR retrotransposons showed that *Gypsy* and *Copia*-type LTRs had a recent and similar burst 0.60-1.00 million years ago (Mya) (Figure 3A), and were enriched far from coding genes (Figure 3B). These results indicated that the burst of *Gypsy*-type LTRs may be the major driving force for the expansion of the tomato genome.

**Figure 3:**
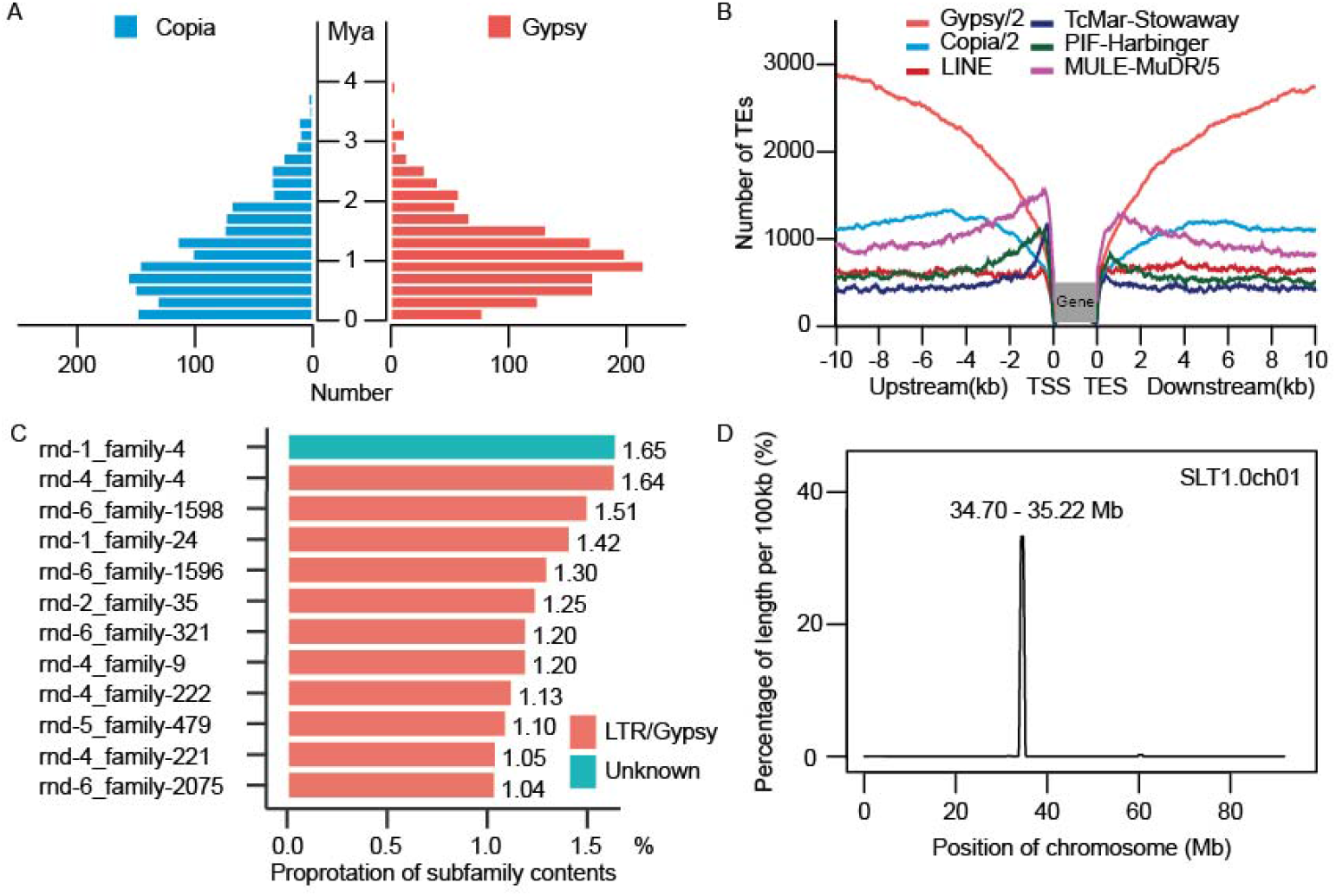
Repetitive sequence analysis. **A** The estimated insertion time of LTR retrotransposons, showing *Gypsy* and *Copia*-type LTRs. **B** Frequencies of transposable elements (TE) in the vicinity of genes. **C** The top 12 TE subfamilies, including 11 *Gypsy* and one *Unknown*-type subfamily. **D** The *Unknown*-type rnd-1_family-4 subfamily was enriched towards the centromere of chromosome 1

To identify the centromere regions, we detected the top 12 TE subfamilies, including 11 *Gypsy* and one unknown-type subfamilies, which together comprised over 15.47% of the genome (Figure 3C). The density of these TE subfamilies along all the chromosomes showed that only the *Unknown*-type rnd-1_family-4 subfamily (1.65% of the genome) was enriched near centromeres but absent from the rest of the genome (Figure 3D and Supp. Figure 2). In addition, we found that 65.21% of the unanchored Contig/Scaffold sequence length comprised highly repetitive regions. Overall, we predicted 12 potential centromeric regions ranging from 1.90 to 6.90 Mb on the 12 chromosomes.

### Comparison of the SLT1.0 and *S. pimpinellifolium* LA2093 genomes

Structural variations (SVs) between wild and cultivated species can cause many phenotypic differences in domestication traits such as fruit weight and quality(Jin et al., 2019). Based on protein homologies between the SLT1.0 and LA2093 genomes, we found that 23,544 genes (68.47%) in the SLT1.0 genome had one-to-one collinear relationships with 23,474 genes (65.64%) in the LA2093 genome (Figure 4A). In addition, genome collinearity analysis showed that syntenic genomic blocks occupied 95.63% of the SLT1.0 genome and 96.67% of the LA2093 genome, respectively. We also identified 6,647 SVs (more than 1 kb in length) between the SLT1.0 and LA2093 genomes, including 3,054 (45.95%) SVs in 2,862 genes (Figure 4B). GO analysis showed that these genes were significantly enriched in the function of oxidation-reduction process, photosynthetic electron transport chain and proton-transporting ATP synthase complex (Supp. Figure 3). We also identified 4,493,889 SNPs and 2,459,597 indels between the two genomes (Figure 4B), including 418,844 SNPs and 245,310 indels located in 29,862 genes. We noted that 45,229 nonsynonymous SNPs resided in 18,178 genes and 9,148 frameshift indels in 1,559 genes, including 7,788 located in domestication regions(Lin et al., 2014). They were significantly enriched in macromolecular complex, pigment metabolic process, nutrient reservoir activity, and intracellular organelle parts (Figure 4C), suggesting these genes may have contributed to disease resistance and fruit traits during tomato domestication.

**Figure 4:**
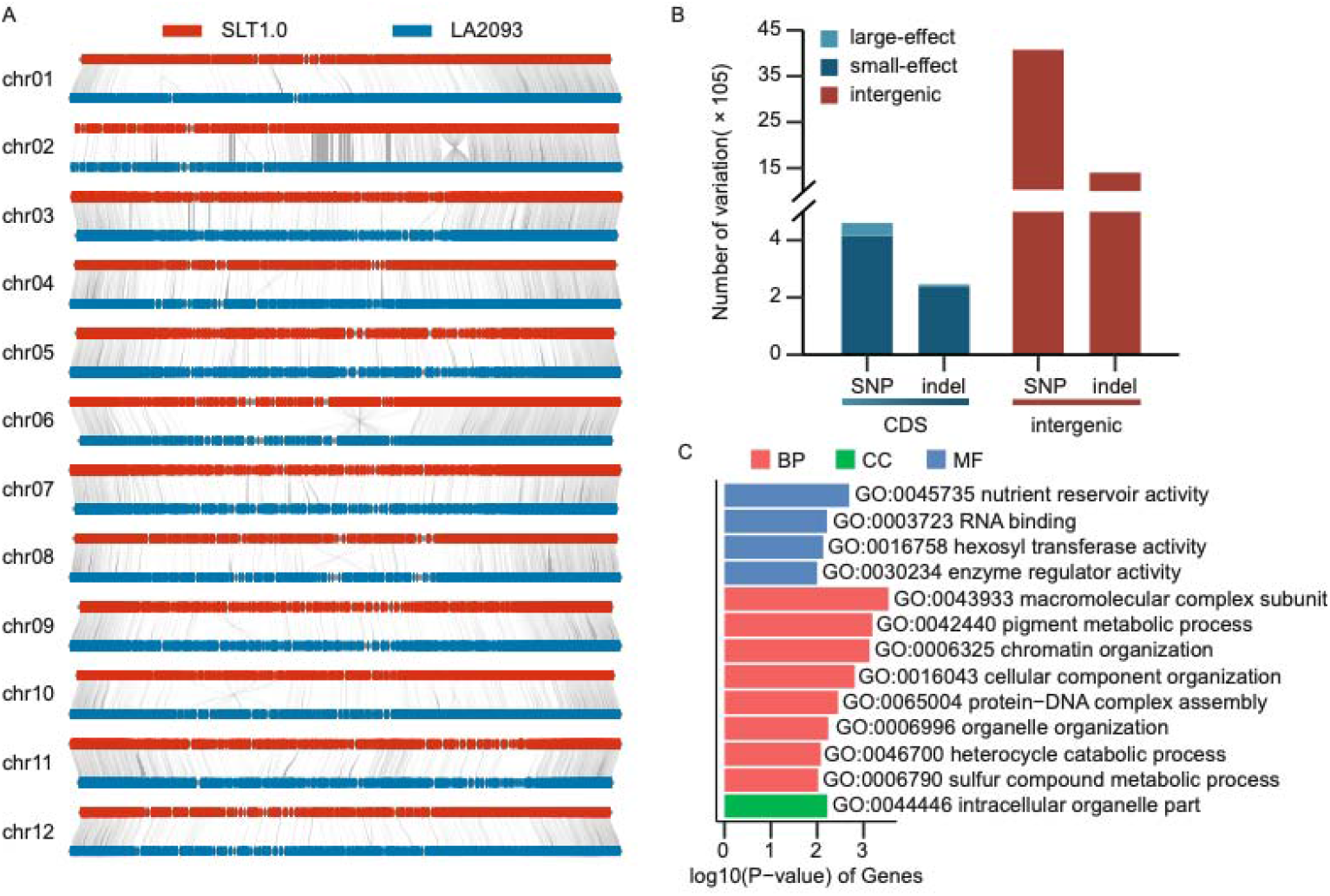
Alignment between the SLT1.0 and *S. pimpinellifolium* LA2093 genomes. **A** The red bar represents the SLT1.0 chromosome, and the blue bar represents the LA2093 chromosome. **B** Numbers of SNPs with nonsynonymous mutations (large-effect), SNPs with synonymous mutations (small-effect), and SNPs in intergenic regions, as well as the number of non-triple (large-effect) indels, triple (small-effect) indels, and indels in intergenic regions. **C** GO terms enriched in genes affected by SNPs and indels selected during domestication

## Conclusion

A highly contiguous and complete genome assembly is a powerful tool for molecular genetic studies of agronomic traits in tomato. In this study, we combined PacBio, BioNano, and Hi-C data to produce the high-quality SLT1.0 tomato genome. The 799.09-Mb assembly had an N50 of 17.83 Mb, and more than 98.94% of its sequences were anchored to 12 chromosomes. The SLT1.0 genome had more repeats were sorted and anchored to chromosomes than the previously released SL4.0 genome. Analysis of repeat subfamilies showed that a specific subfamily, rnd-1_family-4, was found in centromeric regions of the SLT1.0 genome. We could not find a similar reliable repeat family in the SL4.0 genome. Comparative genome analysis revealed that a 2.76-Mb inversion was present on chromosome 2 in SLT1.0 relative to SL4.0 (Figure 2). The inversion was validated by Sanger sequencing and contained no functional genes in adjacent breakpoints, suggesting it is a continuous fragment that has no effect on the SLT1.0 genome. However, we must be cautious and further verify these different fragments between the SLT1.0 and SL4.0 genomes.

Overall, we produced a high-quality tomato genome that will facilitate the molecular dissection of important agronomic traits in tomato. This high-quality genome will be powerful tools for tomato breeding and can deepen our understanding of tomato biology.

## Acknowledgements

This work was supported by the 111 Project (B17043), the Beijing Municipal Education Commission Construction of Beijing Science and Technology Innovation and Service Capacity in Top Subjects (grant CEFF-PXM2019_014207_000032), the National Key Research and Development Program of China (2019YFD1000300) and the National Natural Science Foundation of China (32072571).

## Methods

### Plant materials and sequencing

Plants were grown in the greenhouse in China Agricultural University in Beijing, with a 16 h light/ 8 h dark cycle. A PacBio SMRT library was constructed and sequenced on the PacBio Sequel Ⅱ platform. A Hi-C library was prepared following the Proximo Hi-C plant protocol with HindⅢ as the restriction enzyme for chromatin digestion. The Hi-C libraries were sequenced on the Illumina NovaSeq platform with a read length of 150 bp. For optical mapping, high-molecular-weight DNA was isolated and labeled using a Bionano Saphyr System.

### *De novo* genome assembly

The raw SLT1.0 SMRT reads were corrected and assembled into sequence contigs using CANU with default parameters. The contigs were used for HERA assembly with the corrected SMRT reads. To identify sequence overlaps, all contigs and corrected reads were aligned all-against-all using Minimap2(Li, 2018) and BWA(Li et al., 2009) with default parameters. The HERA-assembled super-contigs were combined with BioNano genome maps to generate hybrid maps using IrysView software (BioNano Genomics) with a minimum length of 150 kb. The resulting contigs were further clustered basing on the Hi-C data using 3D-DNA software(Dudchenko et al., 2017). Pilon(Walker et al., 2014) was used for further error correction.

### Repeat analysis and gene annotation

The integrity of the final genome assembly was assessed in conjunction with BUSCO (v4.1.4)(Simao et al., 2015) using Benchmarking Universal Single-Copy Orthologs. A combination of *de novo* and homology-based methods was used to identify interspersed transposable elements (TEs). A *de novo* repeat library was built using RepeatModeler (v2.0.1)(Bao et al., 2002) and LTR_retriever (v2.9.0)(Ou et al., 2018). Both the *de novo* library and RepBaseRepeatMaskerEdition-20181026, which is the most commonly used repetitive DNA element database, were used to identify TEs with RepeatMasker (v4.1.0)(Graovac et al., 2009).

The RNA-Seq reads from this study were used to predict protein-coding genes in the repeat-masked SLT1.0 genome(The Tomato Genome Consortium, 2012). The cleaned high-quality RNA-Seq reads were aligned to the assembled genome using HISAT2(Kim et al., 2019) with default parameters, and the read alignments were assembled into transcripts using StringTie(Pertea et al., 2015). The complete coding sequences (CDS) were predicted from the assembled transcripts by the PASA pipeline(Haas et al., 2003). The BRAKER(Hoff et al., 2019), GeneMark-ET(Alexandre et al., 2014), and SNAP(Korf, 2004) softwares were performed on *ab initio* gene predictions. Finally, high-confidence gene models were predicted by integrating *ab initio* predictions, transcript mapping, and protein homology evidence with the MAKER pipeline(Cantarel et al., 2008).

### Genome comparisons and SV identification

Genome comparisons between SLT1.0 and SL4.0 and between SLT1.0 and LA2093 were performed via whole-genome alignment using the MUMmer package (v3.23)(Kurtz et al., 2004). The one-to-one alignment blocks were identified using delta-filter program. Then the show-snp tools were used to identify SNPs and indels using uniquely aligned fragments, and the show-diff tool statistics were used to screen for structural variations over 1 kb in length. The SnpEff(Cingolani et al., 2012) software was used to analyze the various SNPs and indel types on the chromosomes.

## Supplementary Legends

**Supplementary Figure 1: Hi-C heatmap of the SLT1.0 genome. The heatmap represents the normalized contact matrix.**

**Supplementary Figure 2: The *Unknown-type* rnd-1_family-4 subfamily was enriched towards the centromere.**

**Supplementary Figure 3: The log_10_(*P*-value) of genes in the domestication region with SV were analyzed by GO enrichment.**

**Supplementary Table 1: Genomic libraries used for genome assembly of Heinz 1706.**

**Supplementary Table 2: Statistics of gene structure among cultivated and wild tomatoes.**

**Supplementary Table 3: Different gene between the SLT1.0 and SL4.0 genomes.**

**Supplementary Table 4: The primer information on chromosome 2 used to analysis the inversion.**

**Supplementary Table 5: Summary of repeats content in the SLT1.0 genome.**

## Notes

### Competing Interest Statement

The authors have declared no competing interest.

## References

Alexandre, L., Burns, P.D. and Mark, B. (2014). Integration of mapped RNA-Seq reads into automatic training of eukaryotic gene finding algorithm. Nucleic Acids Res 119–119.

Bao and Z. (2002). Automated de novo identification of repeat sequence families in sequenced genomes. Genome Res 12, 1269–1276.

Cantarel, B.L., Korf, I., Robb, S.M.C., Parra, G., Ross, E., Moore, B., Holt, C., Alvarado, A.S. and Yandell, M. (2008). MAKER: An easy-to-use annotation pipeline designed for emerging model organism genomes. Genome Res 18, 188–196.

Cingolani, P., Platts, A., Wang, L.L., Coon, M., Nguyen, T., Wang, L., Land, S.J., Lu, X. and Ruden, D.M. (2012). A program for annotating and predicting the effects of single nucleotide polymorphisms, SnpEff. Fly 6, 80–92.

The Tomato Genome Consortium (2012). The tomato genome sequence provides insights into fleshy fruit evolution. Nature 485, 635–641.

Du, H. and Liang, C. (2019). Assembly of chromosome-scale contigs by efficiently resolving repetitive sequences with long reads. Nat Commun 10, 5360.

Du, H., Yu, Y., Ma, Y., Gao, Q., Cao, Y., Chen, Z., Ma, B., Qi, M., Li, Y., Zhao, X., et al. (2017). Sequencing and de novo assembly of a near complete indica rice genome. Nat Commun 8.

Dudchenko, O., Batra, S.S., Omer, A.D., Nyquist, S.K., Hoeger, M., Durand, N.C., Shamim, M.S., Machol, I., Lander, E.S., Aiden, A.P., et al. (2017). De novo assembly of the Aedes aegypti genome using Hi-C yields chromosome-length scaffolds. Science 356, 92–95.

Durand, N.C., Shamim, M.S., Machol, I., Rao, S.S.P., Huntley, M.H., Lander, E.S. and Aiden, E.L. (2016). Juicer orovides a one-click system for analyzing loop-resolution Hi-C experiments. Cell Systems 3, 95–98.

Ellinghaus, D., Kurtz, S. and Willhoeft, U. (2008). LTRharvest, an efficient and flexible software for de novo detection of LTR retrotransposons. BMC Bioinformatics 9, 18.

Giovannucci, E. (1999). Tomatoes, tomato-based products, lycopene, and cancer: Review of the epidemiologic literature. JNCI-J Natl Cancer Inst 91, 317–331.

Graovac, M.T. and Chen, N. (2009). Using RepeatMasker to identify repetitive elements in genomic sequences. Current Protocols in Bioinformatics 25.

Haas, B.J., Delcher, A.L., Mount, S.M., Wortman, J.R., Smith, R.K., Hannick, L.I., Maiti, R., Ronning, C.M., Rusch, D.B., Town, C.D., et al. (2003). Improving the Arabidopsis genome annotation using maximal transcript alignment assemblies. Nucleic Acids Res 31, 5654–5666.

Hoff, K.J., Lomsadze, A., Borodovsky, M. and Stanke, M. (2019), Whole-Genome Annotation with BRAKER. In Gene Prediction: Methods and Protocols, Kollmar, M., 65–95.

Hosmani, P.S., Flores Gonzalez, M., van de Geest, H., Maumus, F., Bakker, L.V., Schijlen, E., van Haarst, J., Cordewener, J., Sanchez Perez, G., Peters, S., et al. (2019). An improved de novo assembly and annotation of the tomato reference genome using single-molecule sequencing, Hi-C proximity ligation and optical maps. bioRxiv 767764.

Jin, L., Zhao, L., Wang, Y., Zhou, R., Song, L., Xu, L., Cui, X., Li, R., Yu, W. and Zhao, T. (2019). Genetic diversity of 324 cultivated tomato germplasm resources using agronomic traits and InDel markers. Euphytica 215, 69.

Kim, D., Paggi, J.M., Park, C., Bennett, C. and Salzberg, S.L. (2019). Graph-based genome alignment and genotyping with HISAT2 and HISAT-genotype. Nat Biotechnol 37, 907–915

Koren, S., Walenz, B.P., Berlin, K., Miller, J.R., Bergman, N.H. and Phillippy, A.M. (2017). Canu: scalable and accurate long-read assembly via adaptive k-mer weighting and repeat separation. Genome Res 27, 722–736.

Korf, I. (2004). Gene finding in novel genomes. BMC Bioinformatics 5, 9.

Kurtz, S., Phillippy, A., Delcher, A.L., Smoot, M., Shumway, M., Antonescu, C. and Salzberg, S.L. (2004). Versatile and open software for comparing large genomes. Genome Biol 5, R12.

Li, H. (2018). Minimap2: pairwise alignment for nucleotide sequences. Bioinformatics 18.

Li, H. and Durbin, R. (2009). Fast and accurate short read alignment with Burrows-Wheeler transform. Bioinformatics 25, 1754–1760.

Li, Q., Li, H., Huang, W., Xu, Y., Zhou, Q., Wang, S., Ruan, J., Huang, S. and Zhang, Z. (2019). A chromosome-scale genome assembly of cucumber (Cucumis sativus L.). GigaScience 8.

Lin, T., Zhu, G., Zhang, J., Xu, X., Yu, Q., Zheng, Z., Zhang, Z., Lun, Y., Li, S., Wang, X., et al. (2014). Genomic analyses provide insights into the history of tomato breeding. Nat Genet 46, 1220–1226.

Meissner, R., Jacobson, Y., Melamed, S., Levyatuv, S., Shalev, G., Ashri, A., Elkind, Y. and Levy, A. (1997). A new model system for tomato genetics. Plant J 12, 1465–1472.

Ou, S. and Jiang, N. (2018). LTR_retriever: A Highly Accurate and Sensitive Program for Identification of Long Terminal Repeat Retrotransposons. Plant Physiology 176, 1410–1422.

Pertea, M., Pertea, G.M., Antonescu, C.M., Chang, T.-C., Mendell, J.T. and Salzberg, S.L. (2015). StringTie enables improved reconstruction of a transcriptome from RNA-seq reads. Nat Biotechnol 33, 290–295.

Shelton, J.M., Coleman, M.C., Hemdon, N., Lu, N., Lam, E.T., Anantharaman, T., Sheth, P. and Brown, S.J. (2015). Tools and pipelines for BioNano data: molecule assembly pipeline and FASTA super scaffolding tool. BMC Genomics 16, 734.

Simao, F.A., Waterhouse, R.M., Ioannidis, P., Kriventseva, E.V. and Zdobnov, E.M. (2015). BUSCO: assessing genome assembly and annotation completeness with single-copy orthologs. Bioinformatics 31, 3210–3212.

Walker, B.J., Abeel, T., Shea, T., Priest, M., Abouelliel, A., Sakthikumar, S., Cuomo, C.A., Zeng, Q., Wortman, J., Young, S.K., et al. (2014). Pilon: an integrated tool for comprehensive microbial variant detection and genome assembly improvement. PLoS One 9.

Xu, Z. and Wang, H. (2007). LTR_FINDER: an efficient tool for the prediction of full-length LTR retrotransposons. Nucleic Acids Res 35, 265–268.

