## Supplementary figures and images for "A high-continuity and annotated tomato reference genome"

### Supp. Figure 1

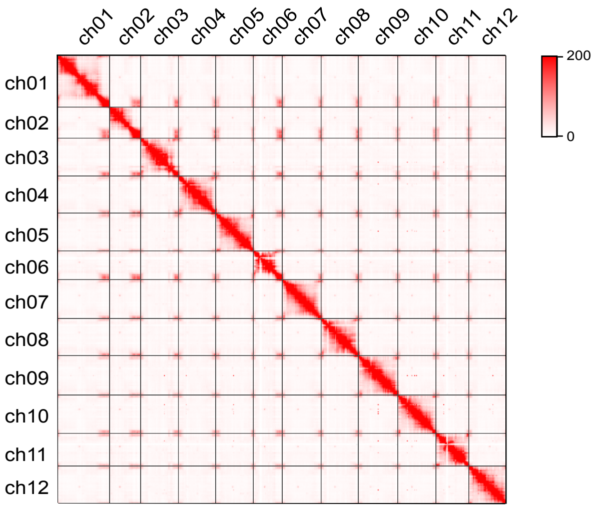

### Supp. Figure 2

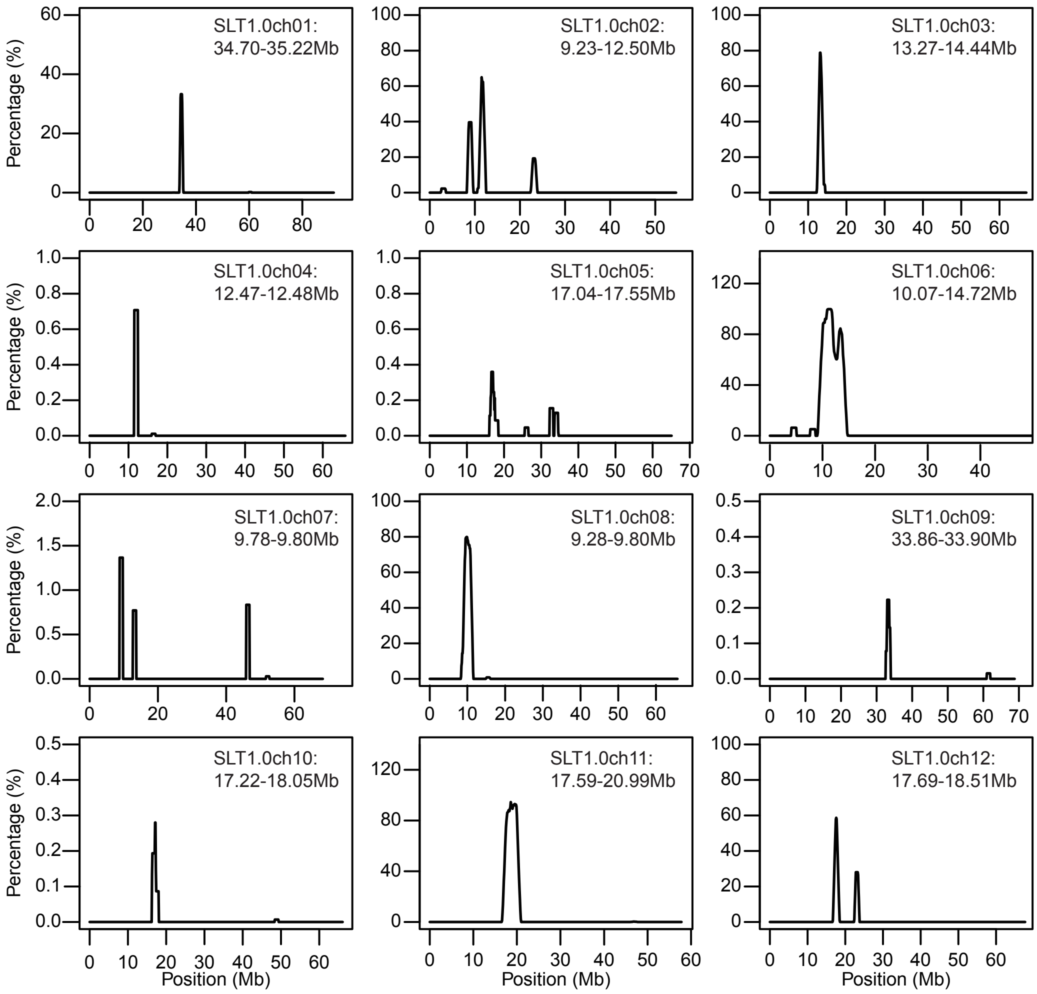

### Supp. Figure 3

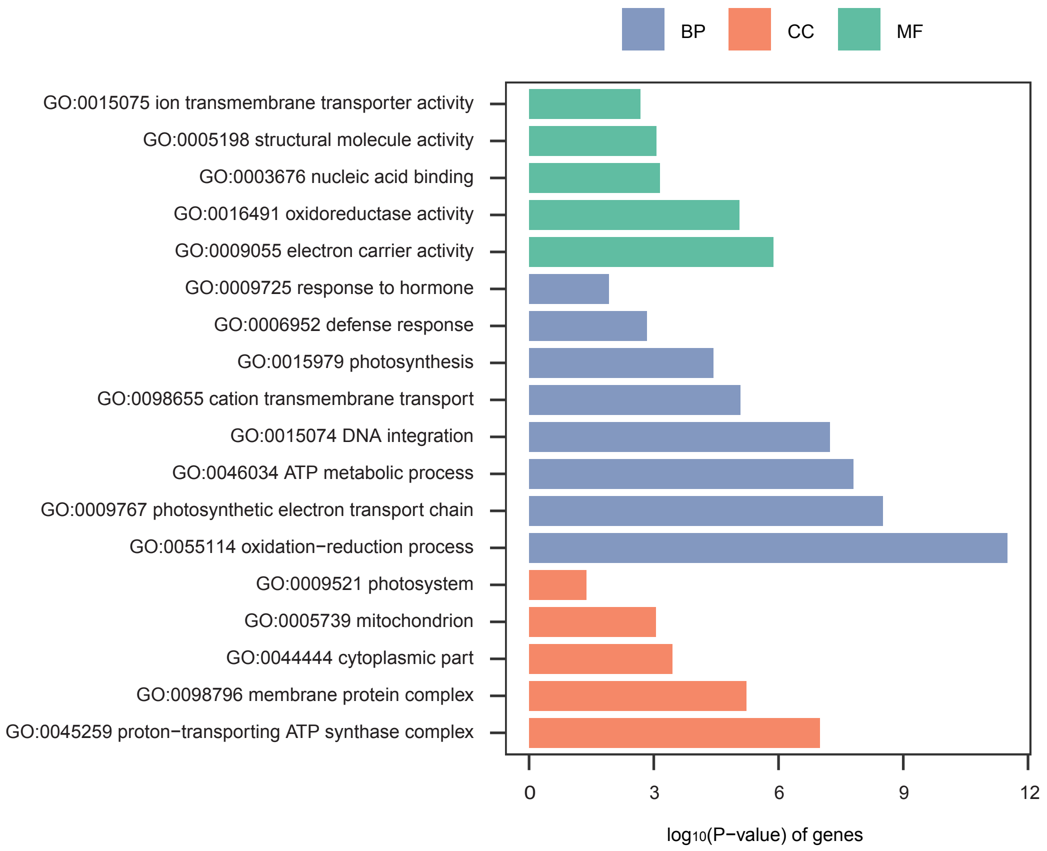
